# Autologous ovarian tissue transplantation and longevity improvement (experimental mice model)

**DOI:** 10.1101/2024.05.06.592661

**Authors:** Nikolai Ruhliada

**Affiliations:** St. Petersburg State Pediatric Medical University, Ministry of Healthcare of the Russian Federation – head of the department of obstetrics and gynecology, Saint-Petersburg, Russian Federation

## Abstract

In the study we showed the improvement in life longevity in mice after step-by step autologous ovarian transplantation. This has proven to be more efficient than “traditional” hormonal replacement therapy. Despite the highest speed and effectiveness of estradiol replacement deficiency in blood by its oral or transdermal use, one did not receive a significant increase in the life longevity in animals and possibly in women (Canderelli R., Leccesse L. et al. 2007). The function of transplanted fragment is usually limited to 6-12 months. This is enough for oncofertility purposes, sometimes, but not for longevity improvement. We performed periodical tissue return (autologous transplantation), containing both cortex and medulla in mice experimental model, that resulted in statistically reliable improvement of longevity. The experimental model we suggested could be projected and to other mammals or humans, as the cortical transplantation gives the same results for reproduction restoration in mice and humans and even on hormone levels normalization, but there is still lack of information about anti-ageing factors containing in ovarian medulla and cortex. That is why we consider that the most important factor for anti-ageing transplantation technology is to preserve both medulla and cortex.

**Summary:** Step-by step autologous ovarian transplantation provides the improvement in life longevity in female mice. This technology could be even more efficient, compared to estrogen hormonal replacement therapy in hormone levels improvement (FSH, estradiol). While menopausal hormonal therapy (MHT) with estrogens doesn’t improve longevity in mice, step-by-step autologous transplantation of ovarian cryopreserved tissue statistically reliably prolongs lifespan in mice.

## Introduction

Ovarian function is one of the important factors in protecting the female body from aging [1, 4]. That is obvious in the experimental model on female mice [1, 3, 5]. Castration leads to a significant decrease in the life expectancy of individuals in comparison with uncastrated animals [4]. The reason for this is both the protective angiogenic function of estrogens, which are manifested by cardio and vascular protective effects, as well as a number of biological substances of the medullary layer of ovarian tissue, which prolong the life expectancy of mice in experiments with whole ovarian transplantation with depleted germ cells and the remaining germinal function [16,20]. In other experiments, authors achieved this by pre-administering to mice VCD (4-vinylcyclohexane diepoxide), which caused complete atrophy of the pool of primordial follicles [2, 6, 24].

One of the mechanisms of aging is the progression of cardiovascular diseases and excessive oxidation, resulted from postmenopausal changes in the endocrine balance. Pathophysiologically persistent metabolic disorder, which is also observed in the aging process, leads to the development of systemic inflammation and disruption of all systems and organs in the body [6]. Circulating lipids and inflammatory mediators interact with each other at several levels, thereby exacerbating the development of chronic diseases, promoting morbidity. Cholesterol and modified lipids can directly activate inflammatory pathways [6, 14].

The attempts of autotransplantation of ovarian tissues has proved the possibility of maintaining both spontaneous ovulatory and endocrine functions in women, widely used in oncofertility cases. However, the time-limited functioning of transplanted grafts (usually no more than a year) meets the goal to save follicles growth, but it is not enough to compensate the hormonal deficiency and slow down the aging processes [3, 6, 31]. At the same time, the exogenous administration of estrogens eliminates only part of the effects that develop during estrogen deficit/castration/in menopause. That is why menopausal hormone therapy (MHT), which is based on exogenic administration of estrogens, does not significantly increase the life expectancy in women, despite the improvement in the quality of life [5,17]. Thus in our experimental model we considered and performed delayed step-by-step autotransplantation every 3 month to support triggering and endocrine effects of autotransplanted fragments.

## Materials and Methods

In order to clarify and compare the role and effects of ovarian factors after ovarian tissue autologous transplantation with MHT, we created a study design to assess the effect of MHT, castration and various options for autotransplantation of ovarian tissues on the life span of mice and the dynamics of transplants hormonal activity [21,24]. For experiment, were selected females of inbred mice (c57BL / 10) weighing 20–25 g at the age of 8 months. Approval and accordance statement : protocols for animal experiments were approved by the Animal Experimental Ethics Committee of the Russian State Pediatric University (Approval no. 1456 /2019) on 12.Feb.2019, in compliance with the National Institutes of Health guidelines for the care and use of laboratory animals. Arrive statement: The present study followed international, national, and/or institutional guidelines for humane animal treatment and complied with relevant legislation from Arrive International guidelines.

None of animals were exposed to/were in the presence of males - separated in age of less than 20 days. Mice of c57BL/10 strain usually become reproductively competent between 45 and 60 days of age and reproductively senescent between 10 and 12 months of age. Ovariectomy at average 250-255 days of age will assure influences the female gonad might have in addition to direct effects of gonadal hormones - thus we make the model of early ageing [23]. Reproductive decline in c57BL/10 mice usually begins with irregular cycles at 8-10 months of age. Surgical procedures are most often conducted in an open field (exteriorizing the tract) and under a dissecting microscope (ovariectomy and transplant procedures). To get the ovarian tissue fragments in diestrus and beginning of proestrus stages we evaluated vaginal smears for exact cycle stage (typical stringy mucous in which are entangled many leucocytes and a few nucleated epithelial cells, none of large cornified cells (squamous) with degenerate nuclei).

Animal experiments were carried out in accordance with the general principles of experiments and the Rules of laboratory practice in the Russian Federation (2003), as well as the provisions of the “European Convention for the Protection of Vertebrate Animals used for experimental and scientific purposes”, Strasbourg, France, 1985. All surgeries were performed under aseptic conditions, subject to mandatory general anesthesia during spontaneous breathing. This protocol describes the procedure used during the technique of ovaries removal and outologous transplantation of ovarian fragments. After weighting and intraperitoneal anesthesia (I.P., 27 ga needle) injection, using a cocktail of ketamine, xylazine and acepromazine (65 mg/kg ketamine, 13 mg/kg xylazine 2.0 mg/kg, acepromazine; cocktail dose - 6.5 ml/kg) we returned the animal to the heated home cage until the anesthesia has taken effect. Once anesthetized, remove the animal from the home cage and place on heated paper towels. Using clippers (#40 blade preferred), remove the hair a few mm lateral to midline on each side, starting just below the ribs and moving distally, preparing a clipped ‘patch’ approximately 2-3 cm square on one side of the prone mouse leaving a strip of hair approximately 1-2 cm covering the midline. Wipe the clipped areas with 70% ethanol and, apply betadine solution to the clipped site with a cotton-tipped swab, apply antibiotic gel for eye protection. Place the animal in prone position on a heated sterile surgical field, and wipe the clipped area with a 70% ethanol. Make a 1.0-1.5 cm paralumbar incision and bluntly dissect the skin from the underlying fascia. In autotransplantation step we do not open fascia, and fix the fragment of defrozen ovary directly to fascia by 4-5 sutures Prolene 7/0 and then close skin with 2 Vycril intradermal sutures 5.0. In case of bilateral ovariectomy we open fascia, locate the adipose tissue that surrounds the ovary in the abdominal cavity and gently extract the fat pad and ovary, identify the ovarian bursa, grasp it with two microsurgical forceps and incise the ovarian bursa opposite the ovarian hilum to expose the ovary. After ovary removal immediately start xenon cryopreservation protocol for ovary, each divided by 6 equal fragments for further autologous return. To get the tissue suspension under dissecting microscope we mince ovarian fragments to pieces, smaller than 0.2 mm^3^ by microblade and microsurgical forceps in saline media (usually 8-10 microfragments for each dose). The mechanically disaggregated tissue suspension is cryopreserved with same xenon slow freezing protocol [18] in a volume of 0.5 ml saline, thus we get the dose for single injection for mice in group 3, further injected subcutaneously via needle gauge 14. After surgical step we place the animal in a heated recovery cage shaded from light and monitor the animal continuously until recovery from anesthesia. Additional meloxicam may be used 24 hours post-operatively if indicated.

All animals were divided in 5 groups - 5 mice in each group, of which 4 groups were surgically sterilized at the age of 8 months and 2 weeks for our study (Table 1). The first group did not receive any therapy after sterilization, while the second group received Divigel (transdermal 17-b ethynilestradiol, OrionPharma) once every 24 hours (applied on ears and shaved areas on the back), starting 30 days after gonads removal. The amount of Divigel (transdermal estradiol preparation) we used for hormonal replacement therapy was estimated 0.5-0.7 mg ethat was equal to 1 g standart recommended dosage for women 65-70 kg weight.

**Table 1.**
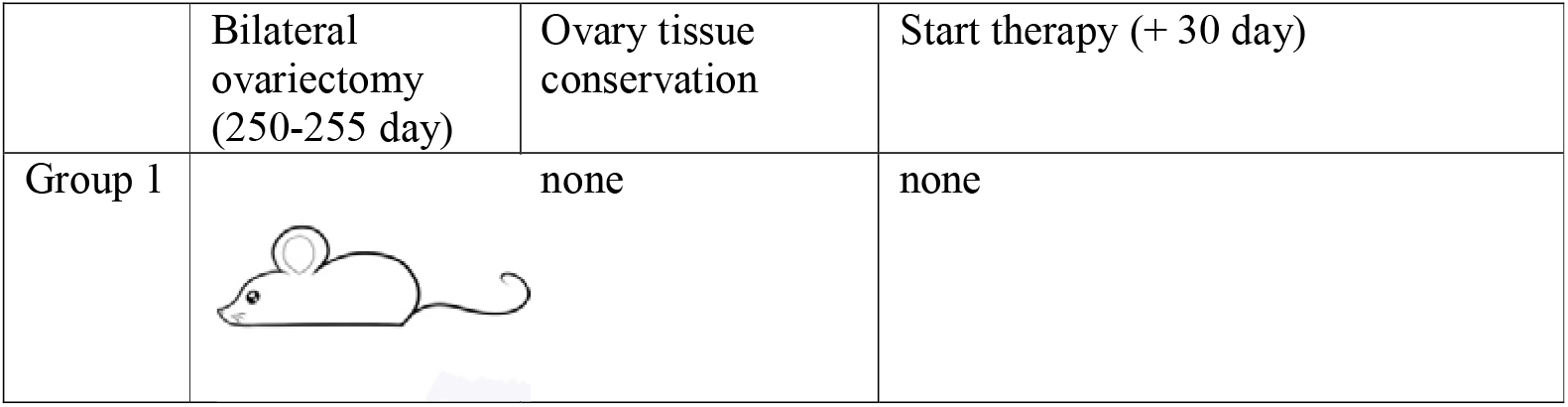
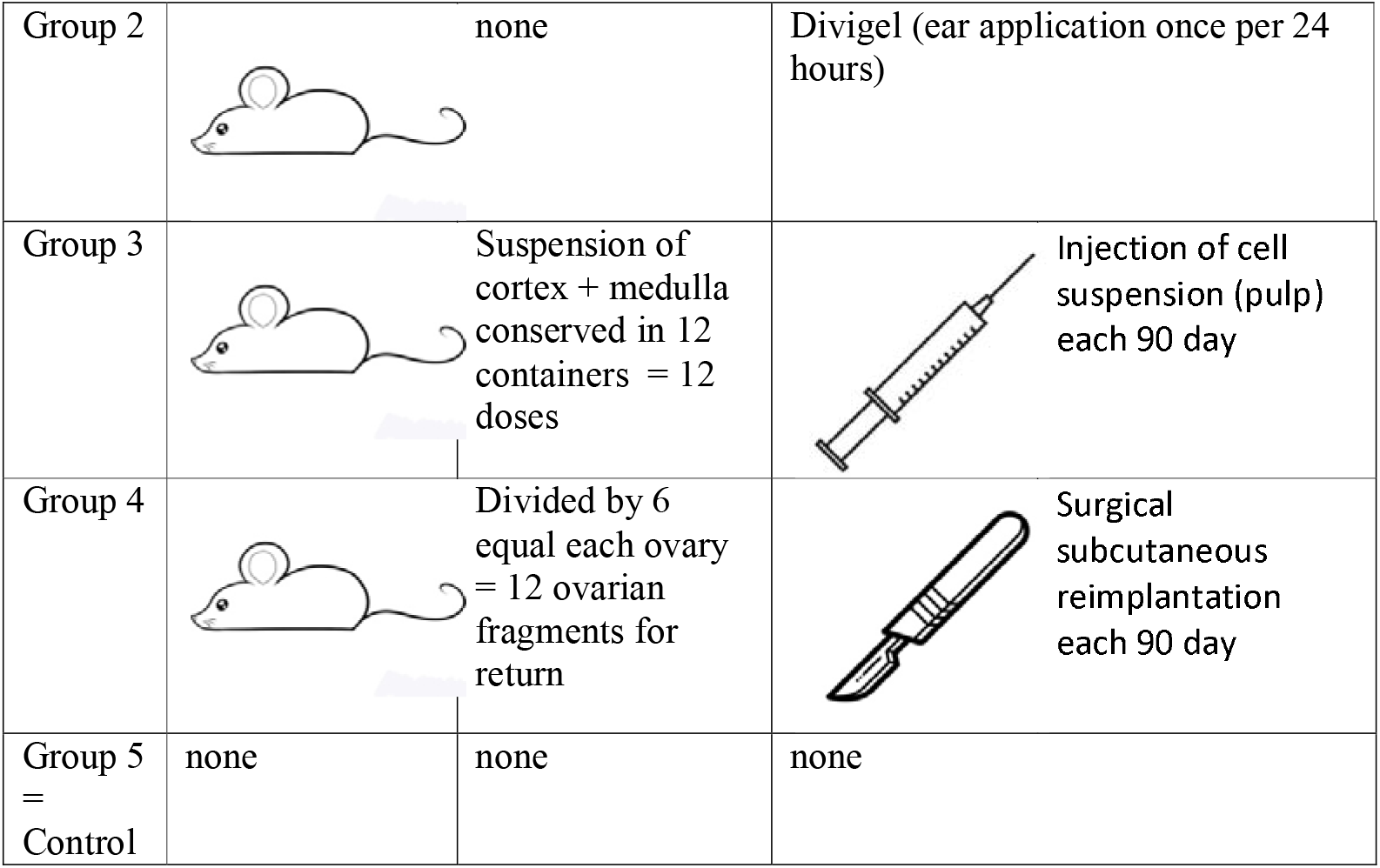
Groups of investigation and research design scheme.

|  | Bilateral<br>ovariectomy<br>(250-255 day) | Ovary tissue<br>conservation | Start therapy (+ 30 day) |
| --- | --- | --- | --- |
| Group 1 |  | none | none |
| Group 2 |  | none | Divigel (ear application once per 24 hours) |
| Group 3 |  | Suspension of cortex + medulla conserved in 12 containers = 12 doses | Injection of cell suspension (pulp) each 90 day |
| Group 4 |  | Divided by 6 equal each ovary = 12 ovarian fragments for return | Surgical subcutaneous reimplantation each 90 day |
| Group 5<br>= Control | none | none | none |

Ovarian tissue was converted into a suspension containing both the cortical and medulla layers, and underwent cryopreservation by slow freezing in xenon medium (purity 99.996%) at a pressure of 1.6 ATM [18] for subsequent subcutaneous injections in group 3, the suspension of each individual was divided into 12 portions for further periodical subcutaneous injections. For mice in group 4, the removed ovaries were divided into 6 parts and also frozen in xenon medium. In the 4th group, 30 days after castration, a quarter of the thawed ovary was reimplanted subcutaneously in the back region (see the protocol above). Third Group – mice, that after castration were administered subcutaneous injections of defrozen ovarian autologous suspension (cortex + medulla) once every 3 months. The first injection is done 30 days after gonads removal. Group 4 - mice, that after castration were subcutaneously transplanted defrozen ovary fragments (approx. 16% = 1/6 ovary volume by one piece) 30 days after gonads removal (cortex + medulla). The first reimplantation in a month after castration, followed by reimplantations each 90 days. Group 5 – mice with no manipulations (control of longevity), consisting of 5 mice, which were not performed any manipulations, and served as a control to assess the comparative life expectancy.

In the studied groups, mice were distributed in separate boxes to exclude the Whitten’s effect [12]. The animals were provided with a periodic similar periodic light in the regime the artificial daylight mode light / dark 16/8, ad libitum nutrition with the same food (protein content 34%, manufactured by Well Plus Trade Vetriebs GmBH, Germany), watered with the same drinking water. The feeding was produced on demand with artificial feed mixtures balanced in terms of proteins, fats and carbohydrates, identical for each group. Feeding of laboratory animals (mice) was carried out with full-feed granulated feed, made in accordance with the standard “Complete feed for laboratory animals” in accordance with the laws of our country [16].

Hormones, including FSH, LH, estradiol, and AMH, were measured at the Clinical Biochemistry Department at our university hospital as normal routine samples. AMH was measured either via the Roche Elecsys assay or the DSL assay (Ansh Lab., Webster, TX, USA).

Statistical processing of the results was performed using the statistical data analysis software package Data were analysed using Stata software (Version 3, Stata Corp). Data are presented as M ± SE, Kaplan-Meier with 99% confidence interval. Significance of differences was evaluated U criteria Mann-Whitney *U* = *nx* · *ny* + (*n*+1)/2 − *T*, the confidence interval was calculated for the probability of p = 0.95 and 0.99 (Table 2).

**Table 2.** Hormones levels in mice on different experiment stages.

|  | Estradiol (pg/ml) | FSH (mMU/ml) | LH (mMU/ml) |
| --- | --- | --- | --- |
| 1 group (castrated) 15 days after gonads removal | 1979±320 | 2,4±1,2 | 2,8±1,3 |
| 1 group (castrated) 30 days after gonads removal | 342±15 | 15±2 | 17±6 |
| 2 group (castrated + Divigel) 30 days after first estradiol application | 3576±445 | 8±1 | 10±2,4 |
| 3 group (castrated + suspension reimplantation) 15 days after 1 injection of tissue suspension | 2345±210 | 16±3 | 14±2 |
| 3 group (castrated + suspension reimplantation) 30 days after 1 injection of tissue suspension | 3245±316 | 0,6±0,2 | 0,4±0,2 |
| 4 group (castrated + 25% ovary volume reimplantation) 15 days after first reimplantation | 1312±450 | 3,7±1,1 | 3,4±0,6 |
| 4 group (castrated + 25% ovary volume reimplantation) 30 days after first reimplantation | 1643±290 | 7,6±3,4 | 11,4±1,1 |
| 4 group (castrated + 25% ovary volume reimplantation) 60 days after first reimplantation | 2854±755 | 1,8±0,4 | 1,1±0,3 |
| 5 group control group of intact mice | 3156±288 | 0,18±0,11 | 0,21±0,08 |

## Results and Discussion

The study of the level of hormones in the blood of mice during the experiment was estimated after castration in group 1, in comparison with the control. A twofold drop in serum estradiol level was observed by 15 days after gonad removal. This rate of decline in estradiol levels corresponds to that for women (1972 and 342 pg / ml) if castrated during surgeries (eg. onco). A gradual increase in FSH was recorded after 2 weeks, but by 30 days it was already statistically significant (15 and 0.18 mIU / ml, p <0.001). In the remaining groups, prior to the start of “return” therapy, the same changes were recorded. Unless the replacement therapy had started. In any way castration has led to equal changes in all groups except control.

But the dynamics of changes in the hormonal profile with the beginning of therapy had its own characteristics in different groups. The higher rate of normalization of estradiol levels and a decrease in FSH and LH we noted with subcutaneous administration (autotransplantation) of a thawed suspension of ovarian tissue. 2 weeks after the first injection, the level of estradiol increased to 2000 pg / ml and higher, and after reimplantation of fragmented ovaries pieces - only to 60th day. This difference is explained by the fact that the speed of revascularization for future functioning of a larger fragment is much lower than with the introduction of a fragmented suspension. Probably, the tissue suspension has more free estradiol, than larger fragments. Nevertheless, in order to compare the effectiveness of one and the second method under identical conditions, we followed the same stages of reimplantation in groups 3 and 4 - once every 90 days (Fig. 1). In all cases, after the natural death of mice at the site of reimplantation of an ovarian fragment or injection of an ovarian suspension, we found signs of vascularization in the subcutaneous layer of cortical and medullary fragments. Thus, the injection of a tissue suspension also leads to the long-term functioning of autologous tissues (Table 3). In both groups 3 and 4 we noted significant decrease in FSH levels in 1 month, as the most reliable marker of ovarian deficiency elimination: 0,6 and 1,8 mMU/ml respectively. Even administration of transdermal estradiol did’t make that fast effect - FSH levels decreased by 8 mMU/ml in 30 days after first estradiol application.

**Table 3.** Mice longevity on different experiment stages.

|  | Average longevity, days | Age at gonads removal, days | Age on the beginning of treatment (transdermal hormones, transplantation autologous ) | Longevity after therapy beginning, days |
| --- | --- | --- | --- | --- |
| 1 group (castrated) | 420 (362-441) | 250-255 | - | - |
| 2 group (castrated + Divigel) | 641 (588-645) | 250-255 | 280-285 | 360 |
| 3 group (castrated + suspension reimplantation) | 930 (796-931)** | 250-255 | 282-285 | 645 |
| 4 group (castrated + 25% ovary volume reimplantation) | 924 (861-952)* | 250-255 | 280-285 | 639 |
| 5– control group | 623 (565-640)*,** | - | - | - |

\*, \*\* - reliable difference $p < 0.001$

The MHT effect of both exogenic estradiol administration and autotransplantation technique led to the best numbers of lifespan. The maximum Kaplan-Meier survival age was in groups 3 and 4 – where the transplantation was performed (930 and 924 mediana days, respectively) (Fig.2). The castration alone, with no MHT resulted in the shortest lifespan – 420 days mediana. Transdermal estradiol andministration improved that parameter to 641 (588-645) days (p<0.001).

Estradiol levels, decreased FSH and LH were not predictors of individuals’ lifespan. Despite the highest speed and effectiveness of replacing estradiol deficiency in the blood by its transdermal use in group 2, we did not receive a significant increase in the life expectancy of animals [7]. It really did not differ significantly from the 5th, control group (641 and 623 days, respectively, p> 0.05), but it was 30-40% lower than when using periodical autotransplantation of the complex of ovarian tissues [8]. Consequently, the periodical transfer of the complex of ovarian tissues (cortex + medulla) allows to increase the life span of individuals, since it “protects” ovarian tissue from physiological exhaustion. The medullary component of the ovary leads to better survival of ovarian tissues during conservation, which is confirmed by Isachenko et al. (2016) [15]. The key role, according to the authors, is played by phosphatidylserine - phospholipid, which plays a signaling function to the activation of apoptosis, and its release from the cortex + medulla complex is significantly lower than with the conservation of only the cortical layer [4, 6, 9].

Our study is consistent with the work of Benedusi V, Martini E, Kallikourdis M (2015), in which the authors showed a decrease in life expectancy with a decrease or complete absence of ovarian activity [4]. Estrogens have an anti-inflammatory effect, thereby reducing the progression of chronic inflammation. However, hormone replacement therapy does not have the desired effect compared to full-fledged ovarian tissue, which leads to the idea that biologically active substances with anti-inflammatory effects are produced in the ovaries.

Indeed, in vitro estrogens act as antioxidants [8]. However, with their low plasma concentration and organism as a whole, they are unlikely to directly have such an effect in vivo [10]. Nevertheless, estrogens cause a pronounced antioxidant effect. After an ovariectomy, the production of H_2_O_2_ by mitochondria grows by 50%. This can be prevented by the introduction of estradiol and transplantation of ovarian tissue [10, 11, 24, 26]. Incubation of cells containing the estrogen receptor MCF7 with estradiol significantly reduces the rate of H_2_O_2_ production. The antioxidant effect of estrogen is mediated by its receptor [8,9,25]. It turned out that the mitochondria of females produce about half of the amount of H_2_O_2_ observed in males. Apparently, this is due to the fact that the active substances of the medullary layer have a higher activity of superoxide dismutase and glutathione peroxidase due to the induction of mitosis-activated protein kinases (MAP) and nuclear transcription factor NF-kB, which trigger the transcription of antioxidant enzymes. Phased replacement of ovarian tissue allows you to activate these processes in which estradiol does not play a major role [27,29]. Our experience indicates the important role of medullary ovarian factors in slowing the aging process of the body and increasing the life expectancy in the experiment [11, 13, 26]. As we showed, the transdermal estrogen supportive therapy in ovarian deficiency improves estrogen levels, but much slower decreases FSH and LH. As well as the best longevity we reached with step-by-step periodic ovarian autotransplantation, thus making “prosthetics” of ovarian function longer, than it is preplanned physiologically (direct correlation between the levels of FSH and lifespan (r=0,98)).

## Conclusion

In the current study, we observed the improvement in life longevity in mice after autologous periodical ovarian transplantation even after primary castration. The model we suggested could be projected and copied to humans as the cortical transplantation gives the same results for reproduction in mice and humans, but there is still lack of information about anti-ageing factors containing in ovarian medullas [25,28,30,31]. That is why we consider the most important factor for anti-ageing transplantation technology is to preserve both medulla cortex, that leads to hormonal triggering effect after autologous tissue return. The improvement in life span could be reached by means of repeated (once 3-9 month) autologous return of ovarian tissue, preserving low FSH levels as a marker of sufficient ovarian activity. The further investigations on dynamics of direct endocrine and triggering effects of transplanted ovarian fragments should be performed prospectively.

## Acknowledgments

The authors declare no competing financial interests.

